# A critical role for human ventromedial frontal lobe in value comparison based on multi-attribute configuration

**DOI:** 10.1101/483719

**Authors:** Gabriel Pelletier, Lesley K. Fellows

## Abstract

Real-life decisions are often between options with multiple value-relevant attributes. Neuroeconomic models propose that the value associated with each attribute is integrated in a global value for each option. However, evidence from patients with ventromedial frontal (VMF) damage argues against a very general role for this region in value integration, suggesting instead that it contributes critically to specific value inference or comparison processes. Here, we tested value-based decision-making between artificial multi-attribute objects in 12 men and women with focal damage to VMF, compared to a healthy control group (N=24) and a control group with frontal lobe damage sparing VMF (N=12). In a ‘configural’ condition, overall object value was predicted by the conjunction of two attributes, while in an ‘elemental’ condition, object value could be assessed by combining the independent values of individual attributes. Patients with VMF damage were impaired in making choices when value was uniquely predicted by the configuration of attributes, but intact when choosing based on elemental attribute-values. This is evidence that VMF is critical for inferring the value of whole objects in multi-attribute choice. These findings have implications for models of value-based choice, and add to emerging views of how this region may interact with medial temporal lobe systems involved in configural object processing and relational memory.

## INTRODUCTION

Everyday decisions are often between options with multiple attributes. For instance, snacks can be characterized on taste, price, and healthiness. Individual attributes might directly predict subjective value: if one craves sweets, chocolate will be valued over peanuts. However, value can also emerge from the interaction of attributes. For example, for those who enjoy “sucré-salé” flavours, the combination of peanuts and chocolate in the same snack would yield a value greater than the sum of the value of each attribute.

Neuroeconomic models propose that subjective value is encoded in a common currency within the ventral prefrontal cortex (Bartra et al., 2013). The overall value of objects composed of multiple value-predictive attributes (e.g. colors and shapes associated with monetary rewards) can be decoded from spatially distributed patterns of BOLD activity in human ventromedial prefrontal cortex (vmPFC) (Kahnt et al., 2011). In non-human primates, activity in the orbitofrontal cortex correlates with the subjective value of juice options varying in taste and quantity (Padoa-Schioppa and Assad, 2006). Studies addressing the mechanisms of value integration have found that functional connectivity between vmPFC and regions representing sensory or semantic attributes increased during valuation (Lim et al., 2013). VmPFC BOLD activity has also been shown to track the difference between cost and benefit signals in amygdala and striatum (Basten et al., 2010). These findings have been taken as evidence that attribute values are integrated into an overall option value representation within vmPFC, subsequently influencing choice (Levy and Glimcher, 2012).

However, direct evidence that vmPFC is required for value integration is lacking. Damage to the vmPFC and adjacent orbitofrontal cortex (together termed ventromedial frontal lobe, VMF) has been shown to change how multi-attribute information is acquired during decision-making (Fellows, 2006), and to affect which attributes influence choice (Xia et al., 2015). When evaluating multi-attribute visual artworks, people with VMF damage differed in how they weighted specific attributes compared to healthy and other frontal-damaged individuals (Vaidya et al., 2017). These observations could be consistent with a deficit in attribute-value integration, as predicted by value integration models. However, these studies of VMF-damaged patients showed that they not only rely on fewer attributes in making value judgments, but also that they systematically neglected specific attributes. This raises the possibility that VMF plays a more specific role in developing value representations of the multi-attribute options that are typical of everyday decision-making. We hypothesized that this region is required for inferring value from the configural relationship between multiple lower-level attributes. By this account, in our opening example, VMF would be required to predict the unique value of peanuts and chocolate together, and perhaps not for simply summing the individual values of each of those attributes alone.

Object processing research has argued for distinctions in the neural encoding of individual attributes and the conjunctions of attributes, with configural processes relying on medial temporal lobe (MTL). Damage to the hippocampus impairs working memory for object-location configurations (Olson et al., 2006) and learning object-outcome associations predicted by attribute configurations (Rudy and Sutherland, 1995). Perirhinal cortex represents complex objects distinct from the combined representations of their parts (Erez et al., 2016) and damage to this region impairs object discrimination based on configurations (Bussey et al., 2005). In contrast, attribute-outcome associations and object discrimination based on individual object parts do not rely on intact MTL. There are strong anatomical connections (Heide et al., 2013) and evidence of functional connectivity (Andrews-Hanna et al., 2014; Eichenbaum, 2017) between MTL and VMF. These regions may interact during decision-making (McCormick et al., 2018, Gluth et al., 2015). Thus, how attributes of complex objects are represented in MTL may be relevant to understanding the role of VMF in assigning value to such objects.

We tested the hypothesis that VMF plays a specific role in inferring value from multi-attribute configurations. We asked whether VMF damage impairs decisions between objects when values were predicted by attribute configurations, by the integration of individual attribute-values, or both.

## METHODS

### Participants

Twenty-four patients with focal frontal lobe damage were recruited through the cognitive neuroscience research database at McGill University. Lesions were characterized with magnetic resonance or computerized tomography imaging, and registered manually to the Montreal Neurological institute standard brain by a neurologist blind to task performance, using MRIcro software (Rorden and Brett, 2000) (available at www.mccauslandcenter.sc.edu/crnl/mricro). Patients were assigned a priori to a group with damage involving VMF, the region of interest in this study (VMF, N=12) or a frontal control group with damage sparing VMF (FC, N=12). Lesion overlap images of the two groups are depicted in **Figure 1**. Twenty-four healthy control participants (HC) matched for age and education were also recruited from a companion healthy control database that draws participants from the Montreal area via community advertisement.

**Figure 1.**
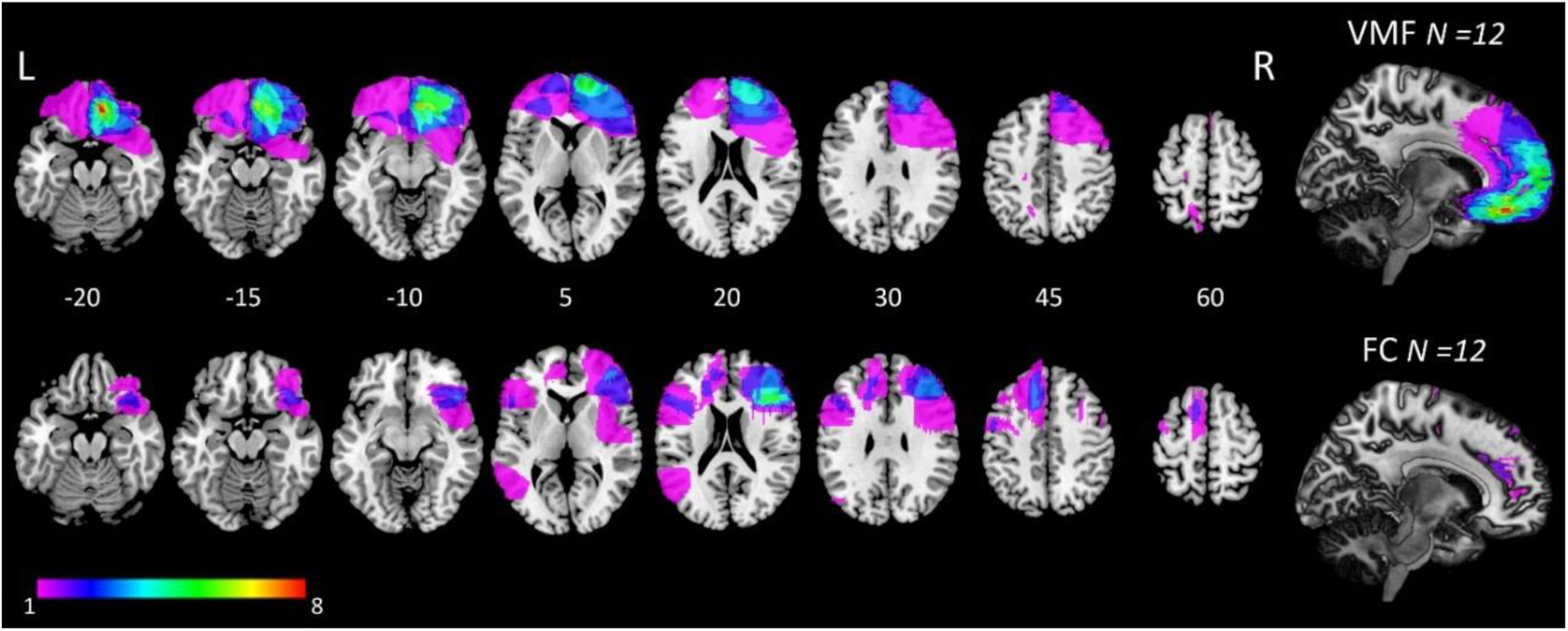
Lesion overlap in the ventromedial frontal (VMF) (top row) and frontal control (FC) (bottom row) groups. Colors indicate extent of lesion overlap, as shown in the Legend. Numbers indicate axial slices by z-coordinate in MNI space.

Damage to the VMF was caused by aneurysm in two cases, hemorrhagic stroke in one case and tumor resection in 9 cases. Damage in the FC group was caused by ischemic stroke in six cases, hemorrhagic stroke in two cases and tumor resection in 4 cases. Nine patients (7 VMF, two FC) were taking one or more psychoactive drugs, most commonly an antidepressant or anticonvulsant. All patients had fixed, circumscribed lesions of at least 6-months duration (mean = 8.3, SD = 4.9 y).

### Neuropsychological screening

Participants with frontal lobe damage completed brief screening tests of working memory (backwards digit span)(Lezak et al., 2012), verbal fluency (Animal, Fluency-F)(Benton et al., 1989), language comprehension (similar to the Token test (De Renzi and Vignolo, 1962)), and incidental memory for faces (Bower and Karlin, 1974).

### Apparatus

All healthy controls and 20 patients were tested in-lab on a desktop computer equipped with a 19-inch monitor. Four participants with frontal damage (three VMF, one FC) were tested at home using a 15-inch laptop computer (Fujitsu). Subjects responded using a standard mouse or keyboard depending on the task. Experiments were programmed in Matlab (version 2014b, The Mathworks, Inc.), using the Psychtoolbox extension (PTB-3)(Brainard, 1997).

### Experimental tasks

Participants made value-based decisions between multi-attribute objects in two conditions, which we term “elemental” and “configural”. They also completed two control tasks to assess object discrimination and memory for single attribute-value associations over a delay. We used novel multi-attribute stimuli developed to study object processing. These pseudo-objects, called fribbles, are composed of a main body and four appendages, each taking one of three possible forms, referred to here as attributes. They were designed to mimic perceptual characteristics of real-world objects (Williams and Simons, 2000; Barry et al., 2014).

To familiarize participants with these novel stimuli and establish that VMF damage did not affect the ability to discriminate fribbles, the session began with a discrimination task that was adapted from a previous study on complex object perception in patients with MTL damage (Barense et al., 2007). The task was divided in two parts. In the first 12 trials, three fribbles were displayed side-by-side; two were identical, one was different. All fribbles had the same main body and three of 4 attributes in common, such that the odd fribble out was distinguished by a single attribute. The participants were asked to select the fribble that was different. Once the response was registered (by a mouse click), feedback was given by surrounding the selected fribble with a green (correct) or red (error) border for 1.5 s before proceeding to the next trial. The second 12 trials followed the same procedure but 5 fribbles were presented: two pairs of identical fribbles and one fribble that could not be paired with any other. Again, participants were instructed to click on the odd fribble out. Importantly, the odd fribble shared all its attributes with at least two other fribbles in the set, such that it could only be identified based on the specific configuration (i.e. conjunction) of two attributes.

The main task had a learning phase followed by a choice phase, for each of two conditions: elemental and configural. Participants learned a total of 6 fribble-value associations in two sets of three by observing the outcomes of mock auctions as fribbles were “sold”, one at a time. Participants were instructed to carefully study the different fribbles and the price for which each was sold. A learning trial started with the presentation of a fribble. After a 2 s delay, the amount for which the item had been sold was presented (**Fig 2**). The fribble and its selling price were displayed until the participant pressed a key to go to the next trial. One learning block included three different fribbles, presented 9 times each in random order for a total of 27 trials. The selling value associated with a fribble on a given trial was randomly drawn from a normal distribution with a standard deviation of 5$; median values are shown in Fig. 2. The fribble associated with each value was randomized, counterbalanced across participants.

**Figure 2.**
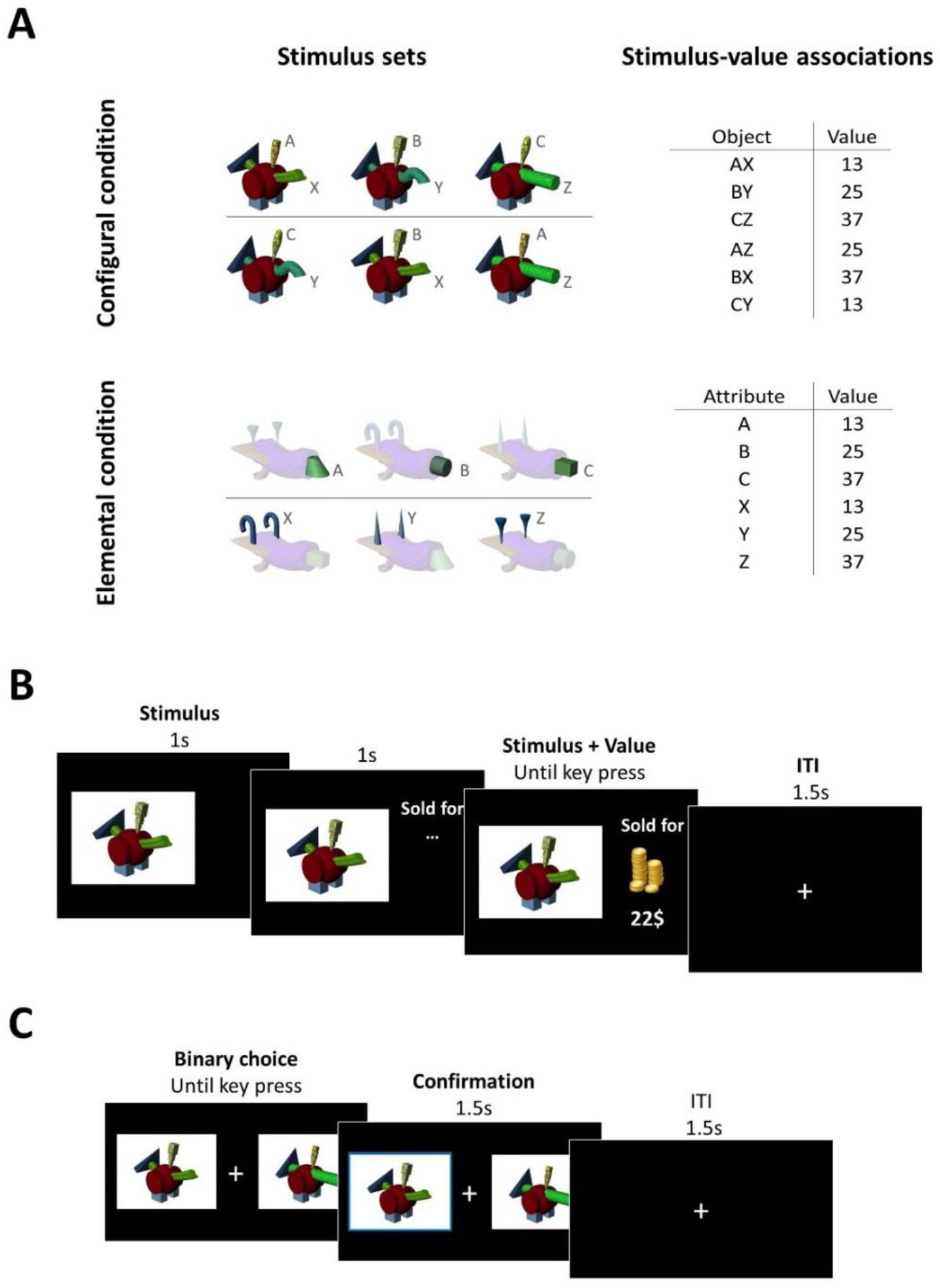
Stimulus sets and experimental paradigm. **A**, Example of stimulus sets and value associations. Stimulus sets were counterbalanced across participants and stimulus-value associations were randomly selected from 6 predefined lists. **B**, Structure of a learning trial. A fribble was displayed for 2 s, then its selling price was presented until a key was pressed to move to the next trial, following a 1.5 s inter-trial interval. **C**, Binary choice trial (for learning probes and final choice phases). These were self-paced: two fribbles were presented on either side of a fixation cross and participants pressed the left or right arrow key to choose which item they wanted in their inventory. Choice was confirmed with a bold border surrounding the chosen object for 1.5 s followed by a 1.5 s inter-trial interval.

After each learning block, learning was assessed with a binary choice probe. On each trial, two fribbles were presented on the screen to the left and right of a central fixation cross (**Fig 2**). Participants were asked to select with the corresponding arrow key which of the two fribbles they thought was worth the most. The response was coded as an error if the less valuable option was chosen, but no feedback was given to the participant. A learning probe block had a maximum of 36 trials (12 repetitions of the three pairs), but was stopped sooner if the learning criterion was violated. Learning blocks and probes were repeated until a criterion of 92% (11/12) correct for each of the three pairs was reached. When criterion was reached with the first set of fribbles, participants were trained on a second set of three following the same procedure.

After learning, participants completed a choice phase which drew upon all six learned associations. On each trial, two fribbles were presented on the screen and participants were instructed to choose which fribble they wanted to have in their inventory (**Fig 2*C***). Participants were told that each fribble they chose would be placed in their inventory, and that this inventory would be sold at the end of the experiment with the proceeds converted into real money (maximum 7$) and added to their base compensation for participation.

In the configural condition, fribbles all had a body and two appendages (attributes) in common. The other two appendages varied such that only the configuration of two attributes predicted the value of the fribble (**Fig 2**). That is, individual attributes did not predict value on their own; their values depended on the other attribute present. During the decision phase, all 15 possible pairs were presented 6 times in random order for a total of 90 trials. In the elemental condition, values were associated with individual appendages (attributes). During the learning phase, the fribble body and irrelevant attributes were masked with a 50% transparent white mask, making the individual value-predictive attribute more salient (**Fig 2**). During the decision phase, stimuli were presented without masks, so all attributes were equally salient, and participants were instructed to take into account everything they had learned about the different parts. The stimulus set included 9 different fribbles (three by three attributes). Thirty-six possible pairs were presented 5 times each, in random order, for a total of 180 trials. Half the trials involved choices between fribbles distinguished by one attribute only (the other attribute being common to the two options), referred to as *single-attribute* trials. Half the trials involved choices between fribbles for which both value-predicting attributes were varied, referred to as *two-attribute* trials. In principle, the more valuable fribble in these trials could be selected by combining (e.g. adding) individual attribute-values, as each attribute was associated with a specific value, regardless of which other attributes were present. All participants completed the configural condition first, to avoid the possibility of an attribute-value ‘task set’ interfering with learning the values of attribute configurations. Stimulus sets were counterbalanced across conditions and participants.

In both the elemental and configural conditions, stimulus-value associations were learned in two different sets before the decision phase. Thus, half the associations called upon in the decision phase were learned more recently than the other half. A control task was therefore included to determine if there were group-wise differences in retaining stimulus-value associations across this delay. This task used a new set of fribbles and was completed after the decision tasks. Three attribute-value associations were trained to criterion, as in the learning phase of the elemental condition. This was followed by an unrelated task (Posner cueing task) lasting approximately 10 minutes, after which memory for the learned associations was probed with a series of binary decisions, identical to the learning probe blocks described above.

### Statistical analysis

Unless otherwise specified, statistical analyses were ran using IBM SPSS Statistics for Windows (version 22). Demographic and neuropsychological screening test variables were compared between patient and control groups using t-tests or Mann-Whitney *U* tests when assumptions for parametric analysis were not met, without correction for multiple comparisons.

Task performance was assessed through accuracy and reaction times. Correct responses were defined as choices of the higher-value fribble in each pair. In the configural condition, each option’s value was the mean value associated with the specific configuration of attributes during training. For the elemental condition, we defined each option’s compound value by summing the mean value of each attribute (learned during training), although any method of combining the two learned values with equal weights would lead to the same stimulus-value ranking. A choice of the option with the lower objectively determined value was coded as an error.

Performance was compared across groups using ANOVA followed by Bonferroni-corrected pairwise comparisons where significant main effects were found. Generalized Estimating Equations (GEEs) were used to analyze the trial-by-trial influence of value on choice behaviour using SAS (version 9.4, SAS Institute Inc., Cary, NC, USA). This analysis is similar to binary logistic regression, but is better suited to modelling repeated measures where outcomes might be correlated within participants, as here. The left minus right option value difference was used as a predictor to model the choice of the left option as a binary outcome.

## RESULTS

### Participant characteristics

Demographic and clinical information is reported in **Table 1**. There was no significant difference in age, years of education or estimated IQ between groups. Patient groups did not differ in lesion volume. The Hospital Anxiety and Depression Scale (HADS) was used to screen for symptoms of anxiety or depression (Zigmond and Snaith, 1983). All but three participants were below the cut-off score for clinically concerning depression or anxiety. One healthy control and two frontal controls scored above the anxiety screening cut-off (HADS-A). One patient in each group scored above the depression screening cut-off (HADS-D). No participant had an active clinically-diagnosed mood disorder, by self-report or chart review. Neuropsychological screening results are shown in **Table 2**. There were no significant differences between patient groups in tests of incidental memory for faces, verbal fluency or language comprehension.

**Table 1.** **Demographic characteristics**

| Group | N | Age (y) | Sex (M/F) | Education (y) | HADS-A | HADS-D | estimated IQ <sup>a</sup> | Lesion volume (cc) |
| --- | --- | --- | --- | --- | --- | --- | --- | --- |
| HC | 24 | 61 (11.6) | 7/17 | 16 (3.0) | 3.8 (3.0) | 2.2 (2.1) | 126 (4) | - |
| VMF | 12 | 57 (10.7) | 5/7 | 14.5 (3.0) | 5.6 (1.7) | 3.9 (3.7) | 120 (8) | 20 (8-192) |
| FC | 12 | 60 (10.7) | 5/7 | 15.1 (2.9) | 5.8 (4.0) | 5.1 (3.6)* | 120 (10) | 24 (5-37) |

All values mean (SD), except sex (count) and lesion volume (median (range)). \**p* < 0.05, Mann-Whitney *U* test compared to healthy controls. HC, healthy controls; VMF, ventromedial frontal damage; FC, frontal controls. HADS, Hospital Anxiety and Depression Scale; A, anxiety; D, depression. ^a^ Not all subjects completed the estimated IQ test.

**Table 2.** **Neuropsychological screening test performance for patient groups**

| Group | Fluency<br>(animals, 60 s) | Fluency<br>(F, 60 s) | Backwards digit<br>span | Incidental memory<br>(accuracy) | Sentence comprehension<br>(accuracy) |
| --- | --- | --- | --- | --- | --- |
| VMF | 10.3 (5.1) | 18.2 (2.9) | 2.6 (0.8) | 0.88 (0.1) | 0.99 (0.02) |
| FC | 10.5 (5.4) | 17.7 (6.7) | 2.9 (1.3) | 0.79 (0.1) | 0.96 (0.07) |

Mean (SD). One VMF and one FC participant did not complete the screening tests.

### Control tasks

Performance of the control tasks assessing the ability to discriminate fribbles and the ability to retain attribute-value associations across a 10-minute delay is presented in **Table 3**. All subjects could discriminate fribbles distinguished by a single attribute or by the conjunction of two attributes. There was no main effect of group on accuracy in the 3-fribble (one-way ANOVA, *F* _(2,45)_ = 0.79, *p* = 0.46) or 5-fribble trials (one-way ANOVA, *F* _(2, 45)_ = 1.47, *p* = 0.24). There was a main effect of group on reaction times in the 3-fribble trials (one-way ANOVA, *F* _(2, 45)_ = 4.82, *p* = 0.01, η^2^ = 0.18). Post hoc comparisons revealed that the FC group was slower than the VMF group (*p* = 0.01), but neither patient group was different from healthy controls (HC-VMF, *p* = 0.26; HC-FC, *p* = 0.22). There was no significant group effect on reaction times in the 5-fribble trials (one-way ANOVA, *F* _(2, 45)_ = 0.61, *p* = 0.55).

**Table 3.** **Control tasks performance for healthy controls and patient groups [mean (SD)]**

| Group | Discrimination-3 fribbles |  | Discrimination-5 fribbles |  | Decision probe after delay |  |
| --- | --- | --- | --- | --- | --- | --- |
|  | Accuracy (%)<br>correct) | RT<br>(ms) | Accuracy (%)<br>correct) | RT<br>(ms) | Accuracy (%)<br>correct) | RT<br>(ms) |
| HC | 99.0 (2.8) | 4940 (1317) | 96.2 (8.1) | 13 248 (6976) | 97.6 (5.6) | 1387 (412) |
| VMF | 99.3 (2.4) | 4165 (667) | 91.0 (13.5) | 11 043 (3378) | 99.3 (2.4) | 1288 (220) |
| FC | 100 (0) | 5755 (1539) | 95.8 (5.6) | 13 913 (8766) | 97.9 (5.2) | 1206 (299) |

Attribute-value associations were retained very well over a 10-minute delay, and there was no effect of group on accuracy (one-way ANOVA, *F* _(2, 45)_ = 0.49, *p* = 0.61) (Table 3)

### Attribute-value learning

All subjects learned the stimulus-value associations to criterion (>92% accuracy) within 4 learning blocks in both conditions. Across groups, more learning blocks were needed to reach criterion in the configural compared to the elemental condition (repeated measure ANOVA, *F* _(1,45)_ = 17.04, *p* < 0.001, η^2^ = 0.28), but there was no significant group by condition interaction (*F* _(2,45)_ = 1.67, *p* = 0.199) (**Table 4**). One healthy control participant was an outlier with respect to reaction times during learning probes (mean 8947 ms and 10765 ms for the configural and elemental condition respectively). After removing this participant from the analysis, we found no main effect of group on reaction times (configural, *F* _(2, 45)_ = 0.79, *p* = 0.46; elemental, *F* _(2, 45)_ = 1.29, *p* = 0.28). The participant with very slow responses nonetheless learned all fribble-value associations to criterion and was included in further analysis.

**Table 4.** **Attribute-value task learning phase performance [mean (SD)]**

|  | Configural Condition |  | Elemental Condition |  |
| --- | --- | --- | --- | --- |
| Group | # of learning blocks | RT (ms) | # of learning blocks | RT (ms) |
| HC | 1.4 (0.5) | 3431 (1654) | 1.2 (0.4) | 2557 (2032) |
| VMF | 1.7 (0.9) | 2504 (664) | 1.0 (0.1) | 1895 (615) |
| FC | 1.7 (0.9) | 3123 (1255) | 1.1 (0.3) | 1647 (702) |

### Multi-attribute value-based choices

Having established that patients with frontal damage could discriminate fribbles, learn elemental and configural value associations, and retain this information across a 10-minute delay as well as healthy controls, we next assessed multi-attribute value-based binary decisions in elemental and configural conditions. The elemental condition involved choices between all possible pairs of fribbles. In principle, half of these trials could be solved by considering the values of single attributes, rather than integrating the values of two attributes, as both options have a value-predicting attribute in common. Trials in which the options differed on both value-predictive attributes (two-attribute trials) require somehow combining the values of two attributes, and were analyzed separately from the single-attribute trials. For the purposes of analysis, we summed the trained attribute-values, but the identical relative value orderings would emerge from averaging these values, or from trading off the values of each individual attribute.

The values learned during training systematically influenced choice in both conditions (**Fig. 3**). Generalized Estimating Equations (GEEs) were used to quantify the extent to which choices were predicted by the difference in option values, trial-by-trial and to test whether this differed by frontal lesion group. Across groups, choices were significantly predicted by the option value difference in both the configural (odds ratio (OR), 2.98; 95% CI, 2.50–3.56; *p* <.0001) and elemental conditions (single-attribute OR, 31.35; 95% CI, 14.52–67.70; p < 0.0001; two-attribute OR, 5.47; 95% CI, 4.54–6.60; p < 0.0001).

**Figure 3.**
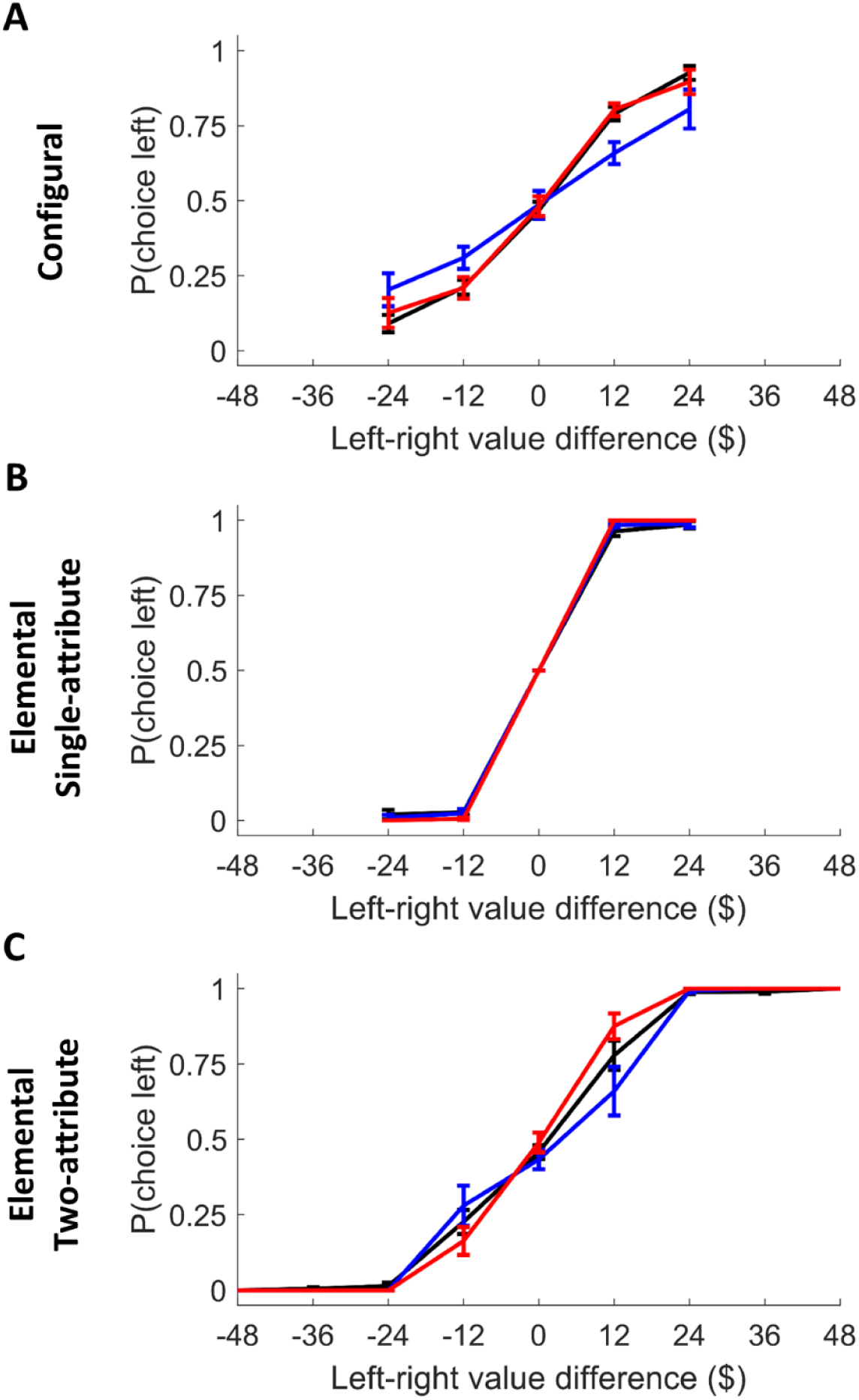
Probability of choosing the left option as a function of the relative value of the left and right options in the configural (**A**) and elemental condition divided in single-attribute (**B**) and two-attribute trials (**C**). Error bars indicate SEM.

The group by value interaction was then added to the model, with healthy controls as the reference group. Compared to healthy controls, the VMF group’s choices in the configural condition were more weakly predicted by option value difference (interaction OR, 0.58; 95% CI, 0.39–0.84; *p* = 0.004). In contrast, the choices of the frontal control group were influenced by option value difference to a similar degree to healthy controls (no significant interaction between group and value; OR, 0.94; 95% CI, 0.61–1.43; *p* = 0.77).

In the elemental condition, the VMF group’s choices were influenced by option value difference to a similar extent as the healthy control group in both the single-attribute (interaction OR, 1.57; 95% CI, 0.34–7.06; *p* = 0.56) and two-attribute trials (interaction OR, 0.93; 95% CI, 0.64–1.37; *p* = 0.73). Value difference was a significantly stronger predictor of choice in the frontal control group compared to the healthy controls in the single-attribute (interaction OR, 17.20; 95% CI, 3.52–84.07; *p* < 0.001) and the two-attribute trials (interaction OR, 1.64; 95% CI, 1.03–2.59; *p* = 0.04). As can be seen in **Figure 3**, the range of value difference was greater in the two-attribute trials of the elemental condition compared to that of the single-attribute trials, and of the configural condition. Because greater value differences are generally associated with easier decisions, and all groups performed at celling at the extreme value differences (**Fig. 3*C***), we restricted the analysis to the two-attribute trials within the same value difference range as the other conditions. We again found no significant difference between the healthy control and the VMF group (interaction OR, 0.87; 95% CI, 0.57–1.34; *p* = 0.53), and value difference was a marginally better predictor of choice in the frontal control group compared to healthy controls (interaction OR, 1.61; 95% CI, 0.99–2.64; *p* = 0.06).

We next asked whether group differences in the influence of value on choice were reflected in significant differences in choice accuracy, defined as the percentage of trials in which the highest value option was chosen. As depicted in **Figure 4*A*,** there was a significant main effect of group on accuracy in the configural condition (one-way ANOVA, *F* _(2, 45)_ = 5.33, *p* = 0.01, η^2^ = 0.19). Post hoc tests with Bonferroni correction for multiple comparisons revealed that the VMF group were less accurate than both the healthy (*p* = 0.01), and frontal controls (*p* = 0.04), whereas frontal controls where not significantly different from healthy controls (*p* = 1.0). There was no significant effect of group on reaction time (one-way ANOVA, *F* _(2, 45)_ = 1.37, *p* = 0.26; **Fig 3*C***) in the configural condition. In contrast, in the elemental condition, there was no significant effect of group on accuracy (one-way ANOVA, *F* _(2, 45)_ = 1.99, *p* = 0.15) or reaction time (*F* _(2, 45)_ = 0.29, *p* = 0.75).

**Figure 4.**
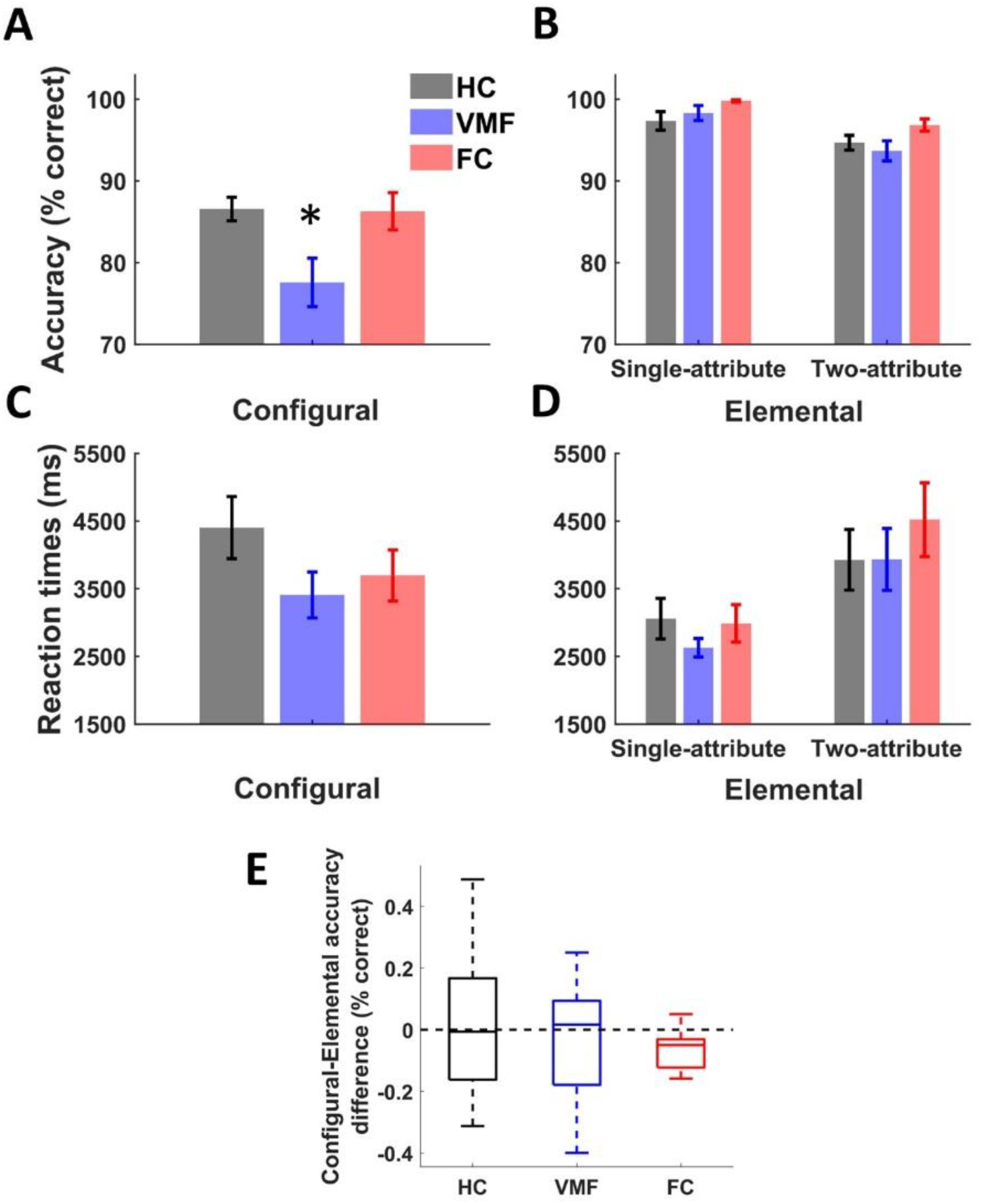
Multi-attribute decision task performance. Choice accuracy for the (**A**) configural and (**B**) elemental condition. Reaction times for the (**C**) configural and (**D**) elemental condition. Error bars indicate SEM, * p < 0.05. (**E**) Difference in accuracy between configural and elemental difficulty-matched trials. Distributions shown with median and quartiles.

Separating the elemental condition in trial types, we found that, across groups, participants made fewer accurate decisions in two-attribute trials compared to single-attribute trials (mixed-measures ANOVA, *F* _(1, 45)_ = 24.0, *p* < 0.0001, η^2^ = 0.35; **Figure 4*B***). There was no main effect of group (*F* _(2, 45)_ = 2.0, *p* = 0.15) on accuracy and no significant interaction between trial type and group (*F* _(2, 45)_ = 0.75, *p* = 0.48). Reaction times were longer in the two-attribute trials (*F* _(1, 45)_ = 46.03, *p* < 0.0001, η^2^ = 0.50) (**Fig 4*D***). Again, there was no significant effect of group (*F* _(2, 45)_ = 1.37, *p* = 0.26) and no interaction between group status and trial type (*F* _(2, 45)_ = 0.29, *p* = 0.75). In summary, VMF damage had no effect on accuracy of decisions between objects based on elemental values, whether for objects distinguished by a single attribute-value, or by two attribute-values.

Finally, we explored whether VMF damage impaired configural more than elemental decisions by directly comparing the two conditions. To make this test as stringent as possible, we selected the trials that were most similar in attribute processing requirements, i.e. the 25 elemental condition trials where both attributes had to be considered to correctly assess value, and the 48 configural trials that were matched with these elemental trials on value difference between options. The difference in accuracy in these trials, for each subject, was then calculated (**Fig. 4*E***), and the relative performance index was compared across groups with non-parametric statistics. We found that the configural-elemental accuracy difference did not differ significantly from 0 in the healthy control (Wilcoxon signed rank test, Z = 0.13, *p* = .90) and the VMF (Z = −0.43, *p* = .67) group, indicating that accuracy was similar between conditions. The frontal control group’s accuracy difference was significantly less than 0 (Z = −2.20, *p* < .05), indicative of lower accuracy in the configural relative to the elemental condition. However, there was no effect of group on accuracy difference (Kruskal-Wallis H test, X^2^(2) = 0.92, *p* = 0.63). This was an exploratory analysis; as evident in **Figure 4*E***, the variance in this subset of trials is high, particularly in the HC and VMF groups, limiting power to detect differences, if present.

## DISCUSSION

We provide evidence that VMF damage impairs value-based decisions between novel multi-attribute objects when overall value is predicted by the configuration of two attributes. This finding was specific to VMF damage: damage to other frontal regions did not impair value-based choices when overall value was predicted by attribute configuration. We did not find evidence that VMF damage impairs decisions between options when individual attributes are independently predictive of value, either when value is assessed based on a single attribute, or when two attribute-values are combined to make an optimal choice.

These findings argue that VMF is involved in assessing the holistic value of multi-attribute objects. This is the first direct evidence that VMF plays a critical role in decisions based on value information provided by the conjunction of individual attributes, each of which is uninformative on its own. The results complement previous work from our lab showing that VMF damage leads to the neglect of some value-predictive information in complex real-world objects (faces, art) (Xia et al., 2015; Vaidya et al., 2017). The current observations raise the possibility that such information may be ‘neglected’ because it relies more heavily on configural processing.

Although the present study was not designed to study value-based learning, it is notable that the learning measures we collected suggest that VMF is not required to learn configural whole-object values through feedback. Configural learning has been shown to rely on the hippocampus (Rudy and Sutherland, 1995), and configural reinforcement learning has been related to the functional coupling between hippocampus and striatum (Duncan et al., 2018). Configural learning in non-human primates is spared after transection of the uncinate fasciculus, disrupting the direct connections between the prefrontal cortex and the temporal lobe (Gutnikov et al., 1997), consistent with our preliminary finding here that VMF damage does not disrupt such learning in humans. As discussed above, VMF becomes critical when object-values must be compared to guide choice. We speculate that there are multiple strategies to solve multi-attribute decisions, and VMF damage may selectively disrupt configural (holistic) strategies, leading to suboptimal choice.

Decisions in the elemental condition could in principle be achieved by option-based or attribute-based strategies, either integrating the attribute values within options and then comparing the compound values, or by comparing individual attribute-values. There is an extensive literature showing that within these broad approaches to such decisions, there are further strategies that may be engaged (e.g. trade-offs, elimination-by-attribute) (Bettman et al., 1998). We cannot address which strategies might have been used here, but prior work on explicitly multi-attribute (elemental) choices where attribute information is presented in tabular format has demonstrated that VMF damage affects these processes, biasing towards simpler, within-option valuation rather than cross-option comparison strategies (Fellows, 2006).

The work clearly shows that VMF damage does not impair learning or choices based on single value-predictive attributes, when those attribute values are explicitly trained. Could VMF also play a role in multi-attribute decisions involving the integration of independent attribute values? Previous fMRI work has shown that activity in VMF tracks the value of items composed of multiple independently value-predictive attributes (Kahnt et al., 2011; Lim et al., 2013). Our findings suggest that intact VMF might not be required for choice in such conditions. However, the analysis directly comparing performance on the subset of trials with the most similar attribute processing and value-difference requirements across conditions did not demonstrate a group by condition interaction. Thus, we cannot exclude that VMF is required when independent attribute-values must be traded-off to make an optimal choice. Further work is needed to provide a definitive test of this possibility.

Participants with VMF damage could readily discriminate between complex objects in a control task that relied on configural object representations, ruling out the possibility that the observed impairment in decisions based on configural value was due to perceptual deficits. Configural object perception has been shown to rely on the perirhinal cortex, a MTL region closely related with the hippocampus. Damage to the perirhinal cortex selectively impairs object discrimination when it relies on attribute configurations (Barense et al., 2007; Bartko et al., 2007). In addition, patterns of BOLD activity in this area relate to complex objects held in working memory but not their separate parts, and are relatively insensitive to viewpoint (Erez et al., 2016), arguing that the perirhinal cortex represents the identity of objects independent of changes in physical characteristics. Our findings suggest that the role of VMF can be understood in similar terms: i.e. VMF is crucial in developing predictive value representations when attributes on their own are ambiguous or separately uninformative. We speculate that perirhinal cortex interactions with VMF may be important for predicting object values based on attribute configurations.

This proposal aligns with other recent efforts to understand how prefrontal cortex and MTL interact, more generally. Synthesizing the common and distinct effects of human hippocampus and VMF damage in a variety of cognitive domains, McCormick and colleagues argued that VMF plays a supervisory role over the hippocampus in initiating and organizing episodic memory retrieval (McCormick et al., 2018). This interpretation mainly stems from studies addressing autobiographical memories and mental scene construction, with so far little causal evidence available with respect to memory for complex objects of the kind commonly featured in neuroeconomics research and everyday decisions. There is some evidence for hippocampal-VMF interactions during value-based decision. One fMRI study found that when imagining the consumption of novel foods composed of two familiar ingredients, both VMF and hippocampus tracked the construction of the compound value (Barron et al., 2013). Interestingly, the results held after controlling for the value of each separate element, indicating that the compound value was distinct from the linear combination of the elements (i.e. configural). Given the role of the MTL in configural processing and our findings here, the putative supervisory role of VMF over MTL in episodic memory retrieval may extend to subjective value construction for multi-attribute objects, particularly when individual attribute-value relationships are insufficiently informative.

An alternative account suggests that VMF (OFC, specifically) encodes the latent (not directly observable) variables of a task to determine the current goals, i.e. representing a cognitive map of task states (Schuck et al., 2016). Task state representations in OFC, as they have been studied so far, are also compatible with a role of VMF in configural decisions. The term ‘configural’ implies that each observable element is not informative alone. Value is instead inferred from the association between elements, with each element being part of multiple associations. Similarly, in Schuck and colleagues (2016), task states were defined by the configuration of task variables, with each unique variable being part of many states. Further work is needed to establish whether these two accounts of the role of VMF, one emerging from computational views of goal-states, the other from complex object processing, reflect the same underlying processes.

This study has limitations. While all patients included in this study had well characterized focal lesions, disruption of underlying white matter tracts (fibers of passage) can affect regions distant from the lesion site (Rudebeck et al., 2013). Converging evidence, especially from non-human primates where more selective lesions are possible, would be helpful in establishing whether effects are caused by white matter disruption, cortical damage, or both. The task also had limited power to assess elemental multi-attribute choices requiring trade-offs, limiting conclusions about whether VMF is also involved under those conditions. Interestingly, we found preliminary evidence that patients with damage affecting other frontal regions had difficulty with such trials, perhaps reflecting the role of lateral and dorsomedial prefrontal cortex in attentional set-shifting (Dias et al., 1996; Vaidya and Fellows, 2016). Further work on the prefrontal mechanisms of individual attribute-value trade-offs in multi-attribute choice is needed. Finally, task order was fixed, because we were most interested in configural processing and wanted to avoid introducing competition between elemental and configural strategies or task sets through the training procedures. For the same reason, we minimized attentional demands in the elemental training condition: all these design choices may be relevant to the pattern of observed effects.

In conclusion, these findings do not support the view that VMF is generically necessary for tracking or comparing value information in a common currency. Under many real-world conditions, the value of complex objects might be better understood as a property emerging from interactions between perception and memory processes, critically relying on VMF when this information is ambiguous and embedded in the relational content between the parts that compose the whole.

## Acknowledgements

This work was supported by the Canadian Institutes of Health Research (Grant MOP 249071) and the Natural Sciences and Engineering Research Council of Canada (RGPIN 2016-06066). We would like to thank Christine Déry for her help with participant recruitment. We would also like to thank the patients and healthy volunteers for their participation.

